# Blocking antibodies against SARS-CoV-2 RBD isolated from a phage display antibody library using a competitive biopanning strategy

**DOI:** 10.1101/2020.04.19.049643

**Authors:** Xin Zeng, Lingfang Li, Jing Lin, Xinlei Li, Bin Liu, Yang Kong, Shunze Zeng, Jianhua Du, Huahong Xiao, Tao Zhang, Shelin Zhang, Jianghai Liu

**Affiliations:** School of Bioscience and Technology, Chengdu Medical College, Chengdu, China; ABLINK Biotech Co., Ltd, Chengdu, China; Kezhi People’s Air-Defense Equipment Co., Ltd, Chengdu, China

## Abstract

The infection of the novel coronavirus SARS-CoV-2 have caused more than 150,000 deaths, but no vaccine or specific therapeutic antibody is currently available. SARS-CoV-2 relies on its spike protein, in particular the receptor binding domain (RBD), to bind human cell receptor angiotensin-converting enzyme 2 (ACE2) for viral entry, and thus targeting RBD holds the promise for preventing SARS-CoV-2 infection. In this work, a competitive biopanning strategy of a phage display antibody library was applied to screen blocking antibodies against RBD. High-affinity antibodies were enriched after the first round using a standard panning process in which RBD-His recombinant protein was immobilized as a bait. At the next two rounds, immobilized ACE2-Fc and free RBD-His proteins were mixed with the enriched phage antibodies. Antibodies binding to RBD at epitopes different from ACE2-binding site were captured by the immobilized ACE2-Fc, forming a “sandwich” complex. Only antibodies competed with ACE2 for recognizing RBD at the same or similar epitopes can bind to the free RBD-His in the supernatant and be subsequently separated by the Ni-NTA magnetic beads. Top 1 lead from the competitive biopanning of a synthetic antibody library, Lib AB1, was produced as the full-length IgG1 format. It was proved to competitively block the binding of RBD to ACE2 protein, and potently inhibit SARS-CoV-2 pseudovirus infection of ACE2-overexpressing Hela cells with IC50 values of 12nM. Nevertheless, top 1 lead from the standard biopanning of Lib AB1, can only bind to RBD *in vitro* but not have the blocking or neutralization activity. Our strategy can efficiently isolate the blocking antibodies of RBD, and it would speed up the discovery of neutralizing antibodies against SARS-CoV-2.

## Introduction

The recent outbreak of a novel coronavirus disease (COVID-19) has emerged from a public health emergency of international concern to global pandemic. Its pathogen, SARS-CoV-2, is a newly identified β-coronavirus. Coronavirus got the family name from the spike(S) protein on the viral particle. The highly glycosylated S protein stays compact in trimeric state, recognizes receptor on the host cell membrane, and then undergoes a series of conformation changes, proteolysis events and membrane fusion to complete viral entry. For vaccines, clinical diagnosis, early prevention and medication, the S protein is the most significant target.

The primary sequences of S protein between severe acute respiratory syndrome coronavirus (SARS-CoV) and SARS-CoV-2 share about 76% identities and 86% similarities, which indicates high possibility of structural homology and the same infection pathway. SARS-CoV and SARS-CoV-2 recognized the same host cell receptor ACE2 for mediating viral entry into host cells. It was reported that SARS-CoV S protein trimer bound to ACE2 at 1:1 in ratio ^[1,2]^. Before infection, RBD of each SARS-CoV S monomer was partially buried in the inactive “down” conformation and not able to bind ACE2 due to steric clash. Once infection started, one monomer turned “up” its RBD to expose enough space to ACE2, inducing further conformational open and loose for proteolysis ^[1,3]^. Atomic-level structural analysis suggested that the spatial interaction and interface between SARS-CoV-2 RBD and ACE2 was mostly in accordance with the SARS-CoV case ^[4]^. Besides, a Cryo-EM structure of SARS-CoV-2 S protein trimer published recently showed that one of the three RBDs was in “up” conformation and naturally exposed the whole interaction interface ^[5]^, while the classic closed symmetric trimer still existed ^[6]^. That might explain why SARS-CoV-2 is much more contagious than SARS-CoV and causing tricky problems worldwide.

No effective cure or vaccine is currently available for COVID-19. Based on structure information above, blocking SARS-CoV-2 RBD is a rational therapeutic approach. Here we developed a competitive biopanning strategy to efficiently isolate blocking antibodies from phage display antibody libraries. Several high-affinity antibodies targeting SARS-CoV-2 RBD and blocking its binding to ACE2 were isolated, and the top 1 lead exhibited a neutralization activity of SARS-CoV-2 pseudotyped VSV infection.

## Materials and Methods

### Recombinant proteins

ACE2-His was purchased from Novoprotein (Shanghai, China). ACE2-hFc and SARS-CoV-2 RBD-His were purchased from Sino Biological (Beijing, China). SARS-CoV-2 RBD-mFc was expressed using ABLINK Biotech’s HEK 293F expression system.

### A Phage display antibody library

A synthetic human Fab antibody library AB1 (LibAB1) was constructed according to a procedure previously described ^[7]^. Human germline immunoglobulin variable segments VH3-30 and VL1-16 were employed as templates, the complementarity-determining regions L3 (CDR-L3) and H3 (CDR-H3) was diversified by the designed mutagenic oligonucleotides. The oligonucleotides were synthesized using the trimer phosphoramidites mix Z (Glen Research) containing codons for 12 amino acids in the following molar ratios: 20% each Y, S &G, 6% each T & A, and 4% each P, H, R, F, W, V & L. The number of positions denoted by Z in CDR-L3 (QQ (Z)n PLT) and -H3 (AR (Z) n (A/G/D/Y) FDY) was varied from 3 to 12 and 8 to 12, respectively. The library size is estimated to be 1×10^12^.

### Standard biopanning

Antibodies against RBD were screened at the first round using a standard biopanning protocol ^[8]^. Briefly, RBD-His was coated on 96-well Maxisorp plates at 4°C overnight. After the coating buffer was decanted, the plate was blocked with 1% polyvinyl alcohol (PVA) at room temperature for 1 hour. 100μl of phage libraries (10^13^pfu/ml) was added per well for 2-hour binding. After washing eight times with PT buffer (0.05% Tween-20 in PBS), bound phages were eluted with 100mM HCl (100μl per well), followed by 5-min incubation. The eluent was transferred into a 1.5ml microfuge tube and neutralized with 1M Tris-HCl (pH 8.0). Half the neutralized phage solution was mixed with 1ml of actively growing E. coli NEB 5-alpha F’ (OD_600_ = 0.8) in 2×YT media containing 10μg/ml tetracycline and incubated at 37°C for 1 hour. 10^10^pfu of M13K07 helper phages were added next and incubated for another 1 hour. The infected bacteria were amplified in 50ml 2×YT medium containing 50μg/ml carbenicillin and 25μg/ml kanamycin, shaking at 200rpm and growing overnight at 37°C. The next day, phages were harvested in precipitant with PEG/NaCl solution and resuspended in PBS buffer for the following rounds of panning.

### Competitive biopanning

After the first round of the standard biopanning, a competitive biopanning protocol that included steps of competitive binding, magnetic separation, elution and amplification (Fig.1), was applied to isolate the epitope-specific antibodies. Briefly, 100μl of ACE2-hFc protein (5μg/ml) was coated on the 96-well Maxisorp plates. The wells were washed and blocked with 1% PVA, and then the mixture of antibody library (10^10^pfu per well) and free RBD-His protein (100ng per well) was added. After a 2-hour competitive binding, the supernatant was transferred into a 1.5ml microfuge tube containing the pre-washed Ni-NTA magnetic beads (GenScript) and incubated on a shaker at room temperature for 1 hour. Beads were collected using the magnetic separation rack and washed by the PT buffer for 8 times. Bound phages were eluted with 100mM HCl (100μl per tube) after 5-min incubation. Beads were collected using the magnetic separation rack, and the supernatant was transfer into a tube for neutralization. Half the neutralized phage solution was mixed with 1ml of actively growing NEB alpha F’ cells and amplified as the standard biopanning protocol. 10μl of the bacterial culture before infection with helper phages was taken, diluted, and grown on the LB plates containing 50μg/ml carbenicillin at 37°C overnight. The single clones were picked up next day for the phage ELISA assay.

**Fig.1.**
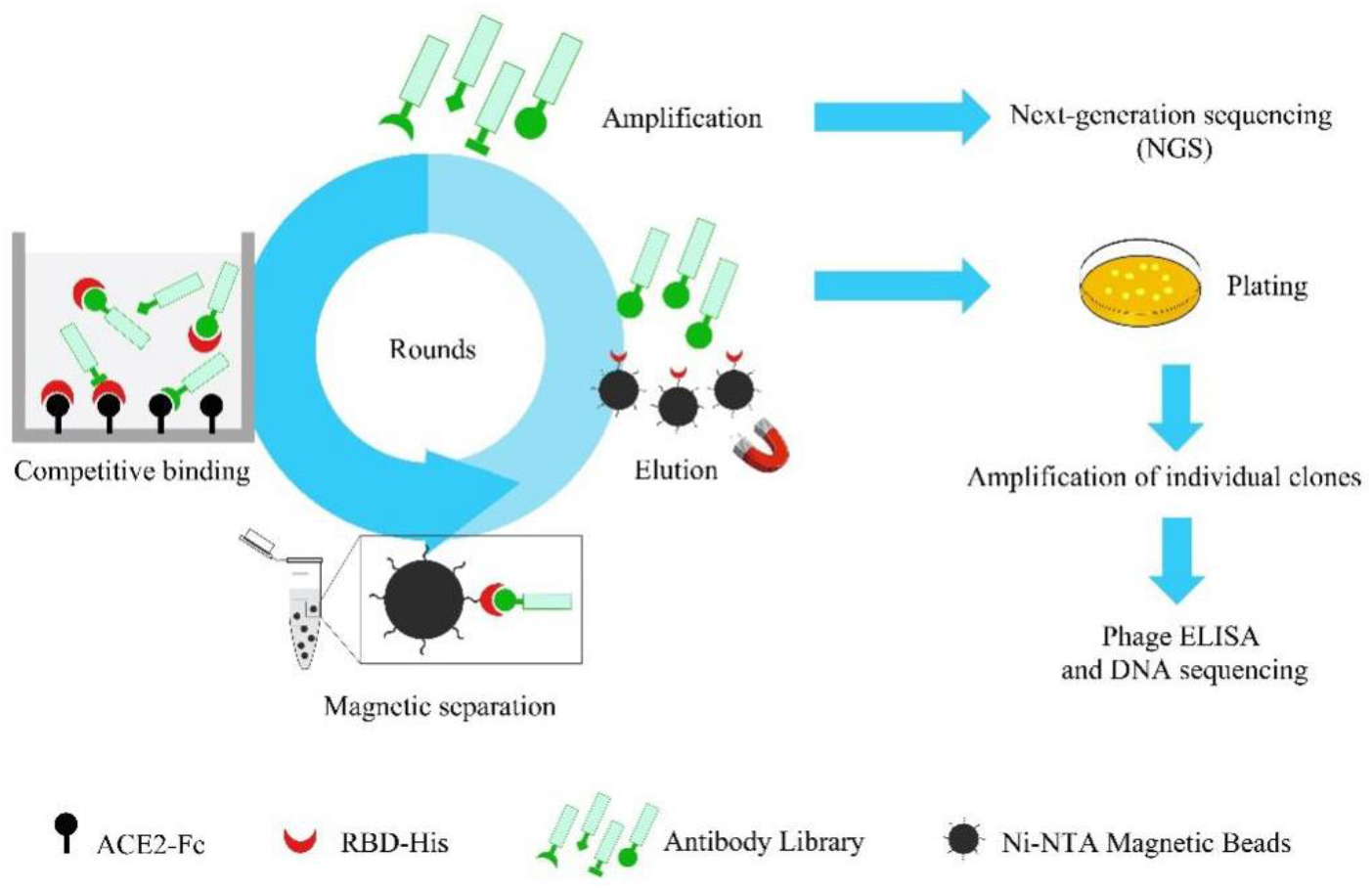
Schematic presentation of a competitive biopanning strategy. A specific binder of target protein was added during the binding step for the selection of blocking antibodies. In this work, the immobilized ACE2-hFc captured RBD-His and the antibodies binding RBD at different epitopes, forming a complex like a “sandwich”. However, when an antibody recognized the same or similar epitopes within RBD as the ACE2 did, it could block RBD-ACE2 interaction. The antibodies would bind to the free RBD-His in the supernatant and be subsequently separated by the Ni-NTA magnetic beads.

### Phage ELISA

Single clones were inoculated into 400μl 2×YT medium containing 50μg/ml carbenicillin, 25μg/ml kanamycin and 10^10^ pfu/ml helper phages in 96-deep-well plates and incubated overnight at 37°C and 250rpm. The plates were centrifuged at 4,000rpm and the supernatant was applied for phage ELISA. The 96-well Maxisorp plates were coated overnight at 4°C with RBD -mFc (1μg/ml, 100μl per well). After blocking with 1% PVA, plates were incubated with 50μl bacterial supernatant containing phages for 2 hours at room temperature. After six times of wash with PT, bound phages were detected using an HRP-conjugated anti-M13 antibody (Sino biological) and tetramethyl benzidine (TMB) as substrate. Absorption at 450nm was measured.

### IgG Expression and Purification

VH and VL of the positive phage were subcloned respectively into the pFUSEss-CHIg-hG1 and pFUSEss-CLIg-hK (Invitrogen). Antibodies were transiently expressed in FreeStyle™ HEK 293‐F cells (Life Technologies) using 293fectin transfection reagent according to manufacturer’s instructions. After transfection, cells were grown in the serum-free medium for an additional 5 days. The supernatant was collected and purified on a MabSelect Protein A column (GE healthcare). Eluted IgG was dialyzed against PBS and stored at −80°C.

### Competitive blocking ELISA

Recombinant human ACE2-His (5μg/ml, 100μl per well) was coated on 96-well Maxisorp plates, followed by a pre-incubated mixture of the anti-RBD antibody titrated into a constant amount of RBD-mFc (1µg/ml). RBD binding to ACE2 was detected using HRP conjugated anti-mouse Fc antibody.

### Pseudotyped virus neutralization assay

The neutralization effects of antibodies on SARS-CoV-2 pseudovirus were performed by the Genscript Inc. (Nanjing, China) under a research service contract. Briefly, 20,000 of the human ACE2-overexpressing Hela monoclonal cells were seeded into each well of a 96-well plate. SARS-CoV-2 pseudovirus and antibodies were incubated at ambient temperature for 1 hour. The mixture was transferred into wells and incubated with cells at 37°C, 5% CO 2 for 24 hours. The culture medium was freshly replaced, and cells were incubated for another 24 hours. The culture medium was removed, and cells were rinsed with PBS. 50µl lysis buffer was added and further incubated at ambient temperature for 40 minutes. 40µl supernatant was transferred to a Sterile Un-Clear 96-well plate with the Bio-Glo luciferase substrate added, and the luminescence signal was measured with EnVision. The dose response curves were plotted with the relative luminescence unit against the antibody concentration. The assay results were processed by Microsoft Office Excel 2013 and GraphPad Prism 6.

## Results

### High-affinity antibodies were identified by the Phage ELISA

After 2 rounds of the competitive biopanning, 48 clones were randomly selected. Their properties of binding to RBD were measured using phage ELISA. Positive binding was defined as an OD450 reading two or more times higher than the negative control (PVA alone). 18 clones showed positive signals (Fig.2). After the DNA sequencing, these clones were summarized into 5 groups of unique antibodies.

**Fig.2.**
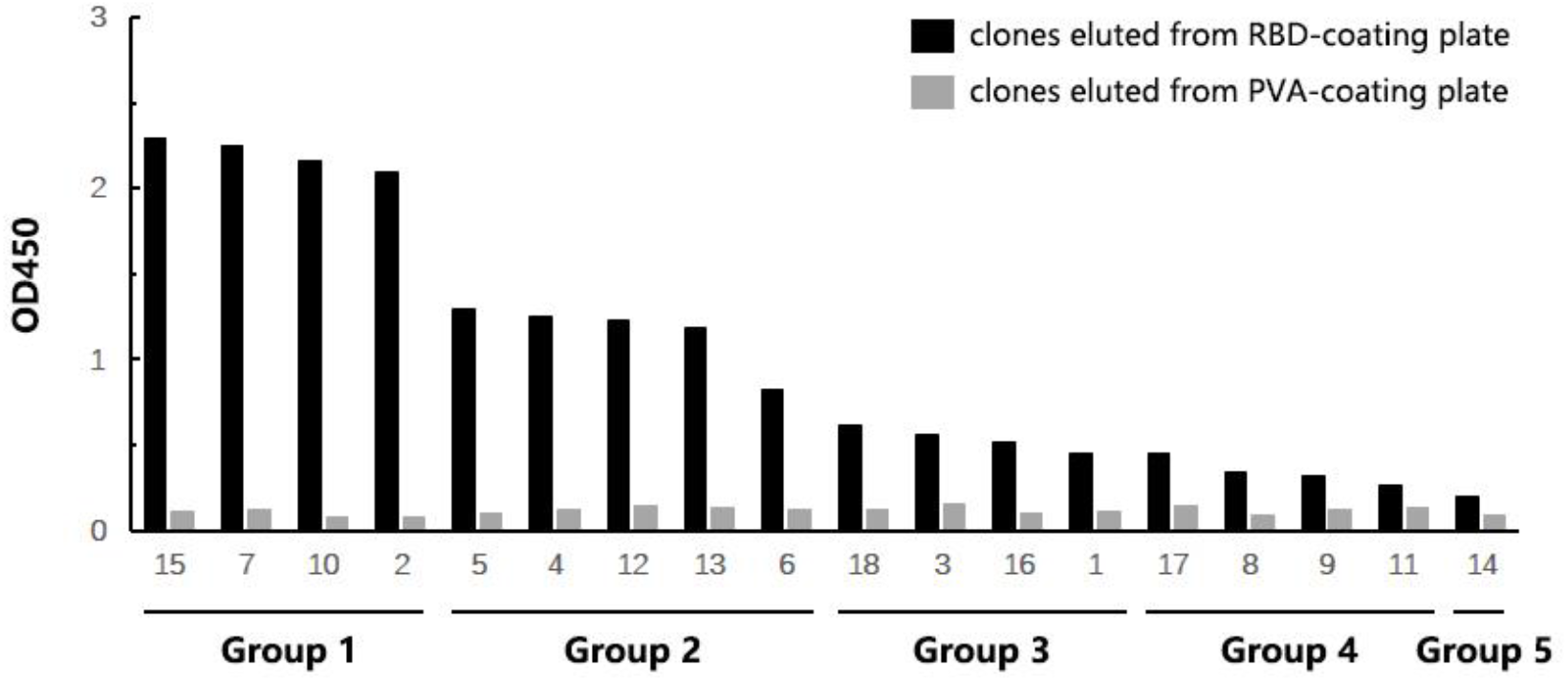
Identification of positive clones to immobilized antigen in competitive manner. Taken OD450 readings as measurement, data of each group fluctuated within 20%. The highest-ranking one was named rRBD-15 in this work.

### Binding and blocking abilities of the top 1 leads against RBD

rRBD-15, the top1 lead with the highest OD450 reading isolated from the competitive biopanning, and rRBD-16, the top 1 lead isolated from the standard biopanning at round 3, were expressed as full-length IgG1 antibodies using the 293F expression system. Their binding and blocking abilities against RBD were compared. Both rRBD-15 and rRBD-16 had high affinities for RBD, with EC50 at 3.8nM and 5.3nM, respectively. Only rRBD-15 blocked the binding of RBD to ACE2 with an IC50 at 3.0nM, while rRBD-16 did not.

### Neutralization abilities of the top 1 leads against SARS-CoV-2 pseudovirus

As a positive control, the recombinant ACE2-hFc (100µg/ml) totally inhibited the infection of ACE2-overexpressing Hela monoclonal cells with SARS-CoV-2 pseudovirus. The antibody rRBD-15 showed a significant neutralization activity against the SARS-CoV-2 pseudovirus with IC50 values of 12.2nM. However, the antibody rRBD-16 had no neutralization effect of the pseudovirus and there were no significant differences between the highest concentration antibody group and the blank group without antibody addition.

## Discussion

RBD-ACE2 interaction initiated viral infection of both SARS-CoV and SARS-CoV-2. Their RBDs share high sequence identities (73%) and structure homology, so the well-established SARS-CoV antibodies were firstly assumed short-cut therapeutic candidates for SARS-CoV-2. However, the real scenario is much more problematic. Several independent peer-reviewed studies as well as preprinted ones have proved that all structurally known SARS-CoV specific antibodies, including S230, 80R, m396 and F26G19, have no cross-reactivity of SARS-CoV-2^[4,5,9]^. These antibodies all compete with ACE2 to bind SARS-CoV RBD, but their epitopes only have limited overlaps of the full ACE2-RBD interface, which could be the reason of lacking cross-reactivity. CR3022 is a special case with 86% conserved key residues in the epitope between SARS-CoV-2 and SARS-CoV. Its cross-reactivity was remarkable, but one site loss of N-glycan results in 1~2 magnitude reduction of binding affinity to SARS-CoV-2 RBD ^[9]^. As in human life, RBD-specific monoclonal antibodies derived from COVID-19 recovered individuals indicated similar patterns of no cross-reactivities with either SAR-CoV or MERS-CoV ^[10]^.

In general, structural and functional analysis suggests that targeting SARS-CoV-2 RBD could be a direct and promising therapeutic strategy, while focusing on previous SARS-CoV antibodies is not very ideal or efficient. No SARS-CoV-2 RBD-specific monoclonal antibody has been reported from human antibody libraries (up to April 17^th^, 2020). In the meantime, SARS-CoV-2 spreads unexpectedly fast around the world, and a new study just shifted its basic reproductive number (R0) from 2.2 to 5.7 ^[11]^. A rapid and effective method of obtaining the SARS-CoV-2 neutralizing antibodies is much required.

Naïve antibody libraries derived from natural immune systems have their capacity limits, while synthetic libraries with higher diversity have more opportunities to isolate binders especially for novel infectious antigens. Compared to a navï e antibody library of 10^8^~10^9^ diversity, a synthetic library with additional artificial randomization on CDRs can reach diversity as high as 10^12^~10^13^. When the recombinant RBD and ACE2 proteins were ready, it took 3 weeks to isolate, produce and verify the antibodies in this study. Using the standard biopanning method, we enriched RBD-specific phages from our synthetic Lib AB1 but not from our navï e antibody libraries (data not shown). Unfortunately, the top1 lead rRBD-16 from the standard biopanning of Lib AB1 couldn’t block the RBD-ACE2 interaction (Fig.4), although it did bind to RBD with an EC50 of 5.3nM (Fig.3).

**Fig.3.**
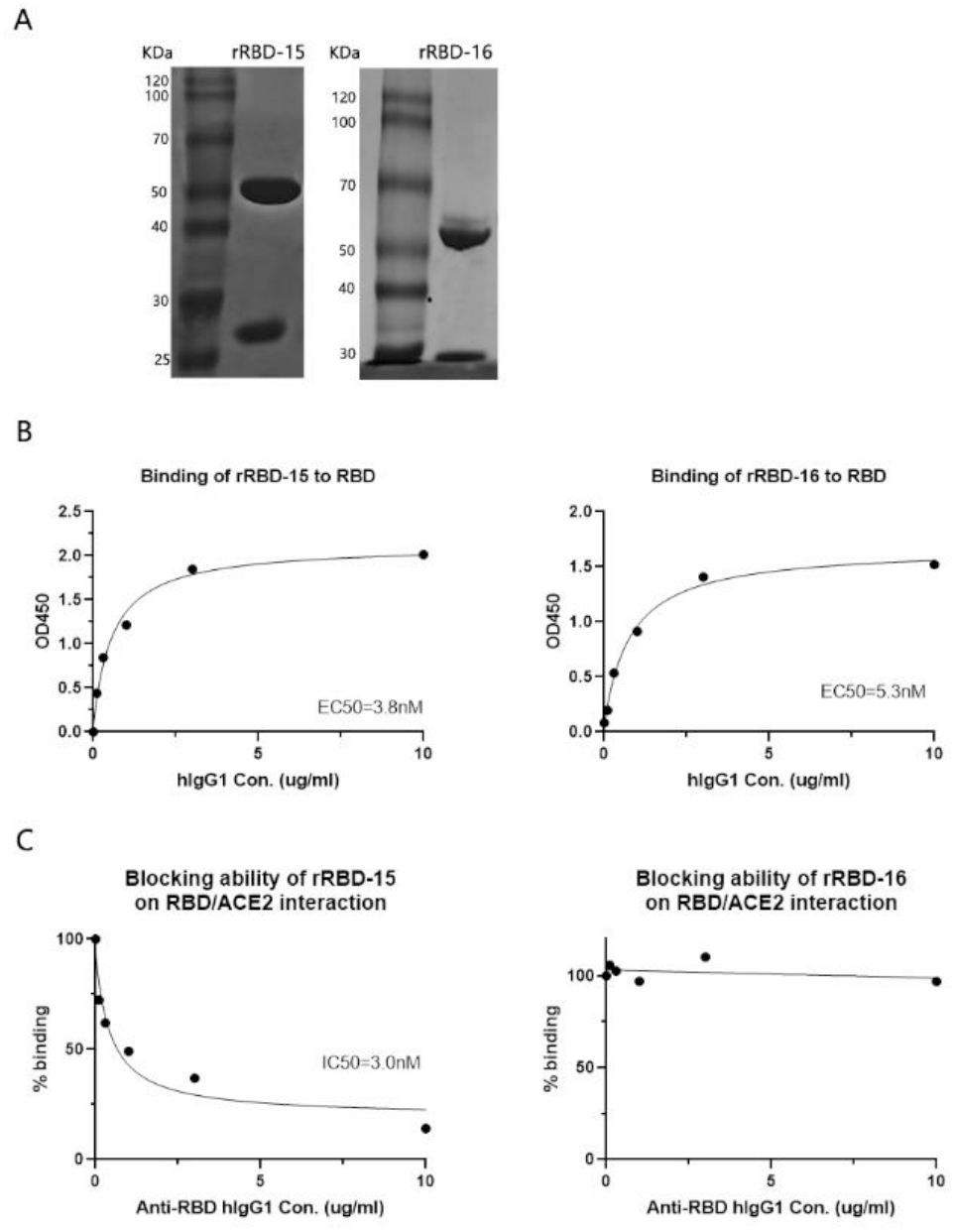
ELISA analysis of the full-length antibodies. (A) Molecular weights of purified soluble IgG antibodies were detected by SDS-PAGE and stained with typan blue. (B) The binding of antibodies to RBD-mFc protein was measured by ELISA. (C) The activities of antibodies blocking the interaction between RBD-mFc and ACE2-His were measured by the competitive blocking ELISA.

**Fig.4.**
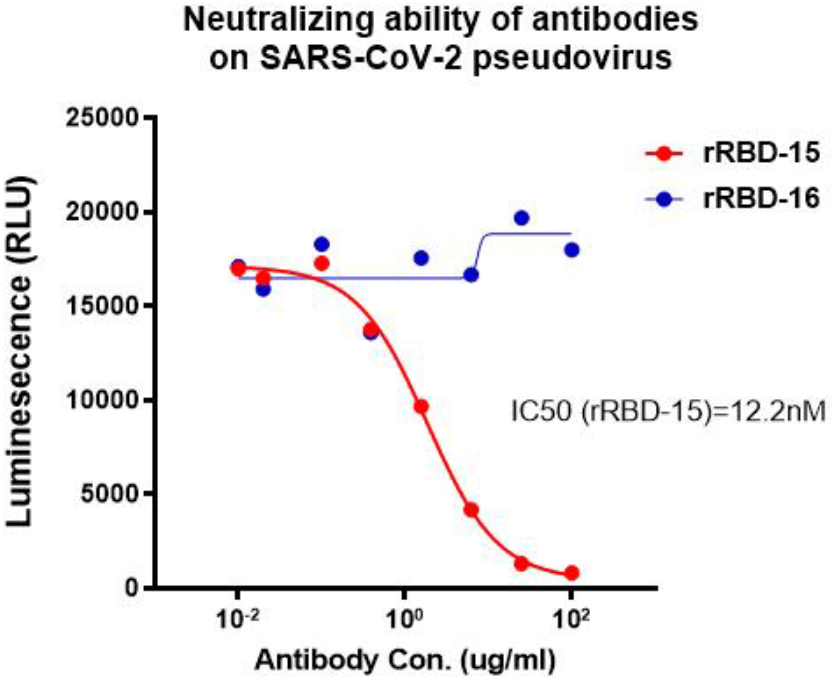
Two SARS-CoV-2 RBD-specific antibodies selected from different strategies showed different neutralization activities. The antibody rRBD-15 competed with ACE2 could neutralize SARS-CoV-2 pseudovirus, but rRBD-16 couldn’t.

The clinical potential and applications of an antibody often depends on its binding epitopes of the target protein. A high-affinity antibody against the target protein can be screened from a phage display antibody library using the standard biopanning process, but its binding epitopes are identified by some extra steps, such as epitope mapping and competitive ELISA. We therefore developed a new competitive biopanning strategy to efficiently isolate isotype-specific antibodies from libraries. As expected, the top1 lead rRBD-15 successfully bind to RBD in compete with ACE2 both in solution and in pseudovirus, and its binding affinity is quite high in 1~10nM differing from measuring methods. In conclusion, our strategic discovery of human monoclonal antibodies against SARS-CoV-2 RBD may fill the blanks of antibody-related pharmaceutical development and shed light on new treatments in need of global health concerns.

## Acknowledgment

We thank Chengdu Zicheng Yibo Biotechnology Co., Ltd for providing the laboratory consumables and bovine serum. This work was supported by Sichuan Science and Technology Program (2018RZ0019), the program of SARS-CoV-2 Protection (CYHX202032, Kezhi People’s Air-Defense Equipment Co., Ltd) and the program of SARS-CoV-2 antibody discovery (JL2020C-01, ABLINK Biotech Co., Ltd).

